# Cognitive modeling informs interpretation of go/no-go task-related neural activations and their links to externalizing psychopathology

**DOI:** 10.1101/614420

**Authors:** Alexander Weigard, Mary Soules, Bailey Ferris, Robert A. Zucker, Chandra Sripada, Mary Heitzeg

## Abstract

**Background:** Individuals with ADHD and other forms of externalizing psychopathology tend to display poor behavioral performance on the go/no-go task, which is thought to reflect deficits in inhibitory control. However, clinical neuroimaging studies using this paradigm have yielded conflicting results, raising basic questions about what the task measures and which aspects of the task relate to clinical outcomes of interest. We aimed to provide a clearer understanding of how neural activations from this paradigm relate to the cognitive mechanisms that underlie performance and the implications of these relationships for clinical research.

**Methods:** 143 emerging adults (ages 18-21) performed the go/no-go task during fMRI scanning. We used the diffusion decision model (DDM), a mathematical modeling approach, to quantify distinct neurocognitive processes that underlie go/no-go performance. We then correlated DDM parameters with brain activation across several standard go/no-go contrasts and assessed relationships of DDM parameters and associated neural measures with clinical ratings.

**Results:** Fronto-parietal activations on correct inhibition trials, which have typically been assumed to isolate neural processes involved in inhibition, were unrelated to either individuals’ response biases or their efficiency of task performance. In contrast, responses to false alarms in brain regions putatively responsible for error monitoring were strongly related to more efficient performance on the task and correlated with externalizing behavior and ADHD symptoms.

**Conclusions:** Our findings cast doubt on conventional interpretations of go/no-go task-related activations as reflecting inhibition functioning. We instead find that error-related contrasts provide clinically-relevant information about neural systems involved in monitoring and optimizing cognitive performance.

## Introduction

The go/no-go task, in which participants are asked to make a motor response following stimulus presentations but to withhold their response after a subset of “no-go” stimuli, is one of the most ubiquitous experimental paradigms in clinical neuroscience. Commonly assumed to index an individual’s ability to inhibit pre-potent or impulsive responses, behavioral metrics from the task, including false alarm (FA) rate and response times, indicate generally poorer task performance in attention-deficit/hyperactivity disorder (ADHD), substance use disorders, and externalizing psychopathology more broadly (1,2,3). Such findings are often cited as evidence in support of the hypothesis that poor inhibitory control is a trans-diagnostic risk factor for externalizing disorders (4,5,6,7,8). In turn, functional magnetic resonance imaging (fMRI) measures of go/no-go task-related neural activations have played a fundamental role in research on the neurodevelopmental mechanisms of inhibitory control (9,10) and on aberrant brain processes in clinical conditions associated with disinhibition (11,12,13,14)

Neuroimaging studies typically focus on several types of contrast images from the go/no-go task when making inferences about inhibitory control. Analyses in which activity during correct rejects (CRs: “no-go” trials where a response is inhibited) is contrasted against activity during “go” trials or a baseline are typically assumed to isolate neural activity related to inhibitory processes (15,16; cf. 17), and tend to reveal right-lateralized activation in prefrontal and parietal structures (18). This approach is based on the subtraction logic that intact inhibitory processes are present during CRs, but not other trials, and hence individuals’ magnitude of activation during CRs should correspond to individual or clinical differences in the integrity of response inhibition (13,16,19,20). In addition, analyses in which activity during FAs is contrasted against that during other trials or baseline may be conducted with the goal of indexing neural activity related to error monitoring. Indeed, such contrasts tend to reveal activation in multiple regions associated with the processing of errors, including the anterior cingulate (ACC), insula and broader fronto-parietal networks (19,20,21,22,23,24).

Despite the go/no-go paradigm’s success in reliably eliciting patterns of neural activation, associations between these activations and clinical outcomes are sometimes difficult to interpret. Across a variety of studies using this paradigm, clinical groups with behavioral disinhibition symptomatology have alternately been found to display reduced (12,19,20,25) or increased (13,26,27,28,29) activation in the brain structures assumed to implement inhibitory processes. Furthermore, significant clinical differences in go/no-go task-related activation are often found in situations where behavioral performance differences are absent (12,14,23,26,28,29). Taken together, these trends in the literature present significant challenges for the conventional assumption that the magnitude of neural responses from go/no-go task contrasts indexes the same underlying construct of inhibitory control that is assumed to be indexed by the task’s behavioral measures.

The current study aims to provide a clearer understanding of how neural activations from the go/no-go paradigm relate to the cognitive mechanisms that underlie task performance and of the implications of these relationships for the study of psychopathology. We utilized data from a large (*N*=143) sample of individuals at risk for externalizing behavior to assess correlations between go/no-go task activations and latent cognitive processes measured by the diffusion decision model (DDM), a well-validated mathematical model of two-choice decision tasks (30,31) that was recently extended to explain performance in the go/no-go paradigm (32,33). The DDM frames the decision of whether to respond or withhold from responding as a noisy evidence accumulation process that drifts between boundaries which represent each of the two possible decision outcomes (Figure 1). Although the process drifts towards, and typically terminates at, the correct boundary (e.g., the “withhold response” boundary on trials with “no-go” stimuli), errors occur when it terminates at the other boundary due to noise. In the DDM framework, several cognitive mechanisms can explain differences in go/no-go task performance: 1) individuals’ general efficiency of evidence accumulation towards the correct choice, as indexed by the drift rate (*v*) parameters, 2) response biases, which are indexed by the start point (*z*) parameter and tend to favor decisions with higher probabilities, such as “go” responses in the go/no-go paradigm (31), 3) caution, as indexed by a parameter for the separation between boundaries (*a*), with lower values indicating a faster but more error-prone decision-making style, and 4) time taken up by processes peripheral to the decision (e.g., perceptual encoding, motor response speed), indexed by a non-decision time (*Ter*) parameter.

**Figure 1.**
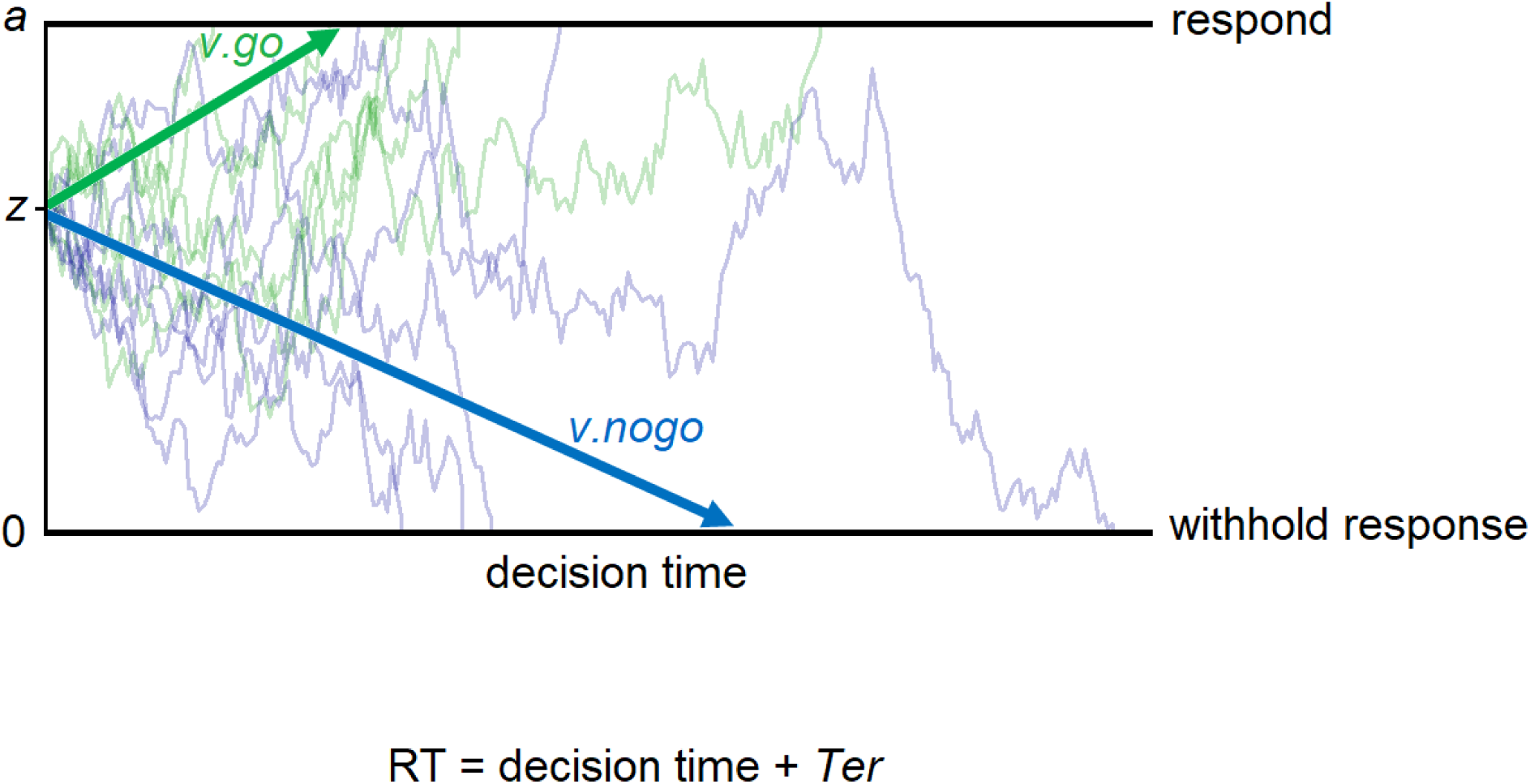
Schematic of the diffusion decision model explanation of the go/no-go task, as outlined by Ratcliff, Huang-Pollock & McKoon (2018) and Huang-Pollock et al. (2017). On each trial, the decision process drifts between an upper boundary for the decision to respond, set at parameter *a*, and a lower (implicit) boundary for the decision to withhold from responding, set at 0. The process begins at the location determined by the start point (*z*) parameter and moves over time according to a stochastic process (similar to a random walk) that generally terminates at the correct decision boundary, but occasionally terminates at the incorrect decision boundary due to noise (i.e., on error trials). The rate at which the process drifts toward the correct boundary (*v*) is estimated separately for trials with “go” stimuli (*v.go*) and trials with “no-go” stimuli (*v.nogo*). RT on a given trial is determined by the “decision time”, which is the amount of time it takes the process to reach one of the two boundaries, and the “non-decision time” (*Ter* parameter), which accounts for time taken up by processes peripheral to the decision (e.g., stimulus encoding, motor response latency). Note that the start point (*z*) is closer to the upper boundary for the decision to respond. This reflects a bias that is expected in a go/no-go task, or any other task where one type of decisional outcome is more likely to be correct than the other (e.g., when “go” stimuli are much more common than “no-go” stimuli), because participants develop an a priori expectation for which outcome is most likely to be correct.

The first goal of the study was to assess whether individual differences in neural activations from common go/no-go task contrasts were correlated with individual differences in one or more of the latent psychological processes indexed by the DDM. The second goal was to assess whether any task-related activation patterns could potentially be interpreted as neural-level indices of mechanistic processes that have meaningful relationships with psychopathology.

## Methods

### Participants

An initial sample of 147 participants, ages 18-21, was recruited from the Michigan Longitudinal Study (MLS) to participate in a neuroimaging study. The MLS is an ongoing prospective study that follows a community sample of families with a history of alcohol use disorder (AUD) and low-risk families from the same neighborhoods (34,35). Participants were excluded from participating in the larger MLS if they displayed signs of fetal alcohol syndrome. Participants were excluded from recruitment into the neuroimaging study if they 1) displayed contraindications to MRI scanning, 2) were left-handed, 3) suffered from a neurological, acute or chronic medical illness, 4) had a personal history of psychosis or schizophrenia, or similar history in first-degree relatives, or 5) were prescribed psychoactive medications in the past 6 months, with the exception of psychostimulants prescribed to treat attention difficulties. Participants using prescribed psychostimulants were asked to discontinue their medication at least 48 hours prior to scanning session. Participants were asked to abstain from using alcohol or illicit substances for 48 hours prior to the study appointment. All study procedures were approved by the University of Michigan’s Institutional Review Board, and all participants provided written informed consent.

Three participants were excluded because their behavioral data did not meet quality control criteria for model-based analyses (at least 200 available trials and overall accuracy rate >.55) and one participant was excluded because they did not commit any FAs, precluding analysis of FA contrasts. This left a final sample of 143 participants (87 males), whose demographic information is displayed in Table 1.

**Table 1.** Characteristics of the sample and descriptive statistics for cognitive and behavioral measures. For continuous variables, numbers indicate the mean with standard deviations in parentheses. ASR summary statistics do not include the 5 subjects with missing data on these measures (all were male). FH-AUD = family history of alcohol use disorder (either parent)

|  |  |
| --- | --- |
| Gender (Male/Female) | 87/56 |
| Age at scan | 19.66 (1.22) |
| Race/Ethnicity |  |
| Caucasian | 128 |
| Hispanic/Latino | 5 |
| African American | 6 |
| Other/bi-racial | 4 |
| FH-AUD (Positive/Negative/Unknown) | 108/34/1 |
| ASR Externalizing – raw score | 9.43 (7.90) |
| ASR Externalizing – T score | 48.98 (10.13) |
| ASR ADHD Symptoms – raw score | 5.02 (4.26) |
| ASR ADHD Symptoms – T score | 54.33 (6.56) |
| Drift rate for “go” stimuli (*v.go*) | 2.77 (1.03) |
| Drift rate for “no-go” stimuli (*v.nogo*)* | 1.90 (1.88) |
| Average drift rate for all stimuli (*v.avg*) | 2.34 (0.88) |
| Boundary separation (*a*) | 1.00 (0.21) |
| Non-decision time (*Ter*)** | .317 (.032) |
| Response bias (*z*) | 0.62 (0.07) |
| False alarm (FA) rate | 0.28 (0.16) |
| Hit RT mean (MRT)** | .448 (.061) |
| Hit RT standard deviation (SDRT)** | .105 (.055) |

\*=Multiplied by -1 for comparability to *v.go*; \*\*=seconds

### Go/No-Go Task

Participants completed an event-related go/no-go task (10) during fMRI data collection in which they were presented with a series of letters for 500ms at a time (interstimulus fixation interval of 3500ms) and asked to press a button for every letter other than “X” (“go” trials) but to withhold their response on trials where an “X” was presented (“no-go” trials). Participants completed 5 182-second imaging runs of 49 trials each, for a total of 245 trials, 60 (25%) of which were “no-go” trials. Neuroimaging data acquisition parameters and sequences (36) are reported in Supplemental Materials.

### Psychopathology Measures

Participants completed the neuroimaging study in between MLS data collection waves. At waves 6-8 (ages 18-24), participants filled out the Adult Self Report (ASR: 37), a questionnaire which assesses levels of internalizing and externalizing psychopathology and symptoms of DSM-IV syndromes. For each participant in the current study, the administration of the ASR closest in time to the scan was identified (mean days between ASR administration and scan=421, SD=401), and raw scores from this measure were used. Five participants (all male) did not have ASR data available (i.e., they did not complete waves 6-8). Therefore, they were not included in analyses that involved prediction of psychopathology.

### DDM Analysis

The go/no-go version of the DDM outlined by (32) and (33), which assumes that “no-go” decisions are made when the decision process reaches an implicit (non-response) boundary, was fit to data in R (38) using the chi-square minimization procedure described in these studies. Functions from the R package *rtdists* (39) were used to calculate chi-square values and simulate model-predicted data to assess fit. As between-trial variability parameters in the model are difficult to estimate without massive numbers of trials (40) and “simple” versions of the DDM, without these parameters, provide estimates of the main model parameters that appear to be comparably reliable and informative to those of the “full” DDM (41), only the main DDM parameters were estimated: drift rates for “go” and “no-go” stimuli, (*v.go*, *v.nogo*), starting point (*z*), boundary separation (*a*) and non-decision time (*Ter*). The upper boundary was assumed to trigger responses, while the lower boundary was assumed to be the non-response boundary. Start point (*z*) was estimated as a proportion of *a*, and *z* values greater than .50 therefore indicate the expected bias towards upper response boundary. Additional information on model specification, estimation, and checks of model fit is available in Supplemental Materials.

### fMRI Data Pre-processing

Functional images were first reconstructed using an iterative algorithm (42). Head motion was corrected with realignment using FSL 5.0.2.2 tools (FMRIB, Oxford, United Kingdom) and runs were excluded if they exceeded 3 mm translation or 3° rotation in any direction during the run. Remaining pre-processing steps, which were carried out using Statistical Parametric Mapping 8 (SPM8: Wellcome Institute of Cognitive Neurology, London, United Kingdom), included spatial normalization to standard space as defined by the Montreal Neurological Institute template, resampling to 2×2×2mm voxels, and spatial smoothing with a 6mm full-width half-maximum Gaussian kernel.

### fMRI Analyses and ROI Selection

A general linear model (GLM) was fit at the individual level with three main regressors convolved with the hemodynamic response function: 1) correct go trials (hereafter referred to as the GO condition), 2) correct rejection (CR) no-go trials, and 3) false alarm (FA) no-go trials.

Motion parameters from earlier realignment and average white matter signal intensity for each volume were also included as nuisance regressors. Following individual-level analyses, four group-level contrasts were conducted: CR>GO, FA>GO, CR>FA, and FA>CR. Clusters from the resulting statistical maps were determined to be statistically significant if they met a cluster-level FWE-corrected threshold of .001. From these thresholded maps, we selected a smaller number of discrete clusters as regions of interest (ROIs) based on previous research (detailed below in Results). Average individual-level contrast parameter estimates were then extracted from each ROI using MarsBaR (43).

### Data Analytic Plan

Following selection of ROIs, we conducted three main analyses to accomplish our goals. Analyses were conducted within R or JASP (44), an open-source statistical package which allows frequentist and Bayesian versions of common statistical tests to be easily implemented. First, with the goal of identifying relationships between parameters of the DDM and task-related neural activations, we conducted Bayesian correlation analyses in JASP between each DDM parameter and activation estimates from each ROI. Bayesian analyses allow estimation of 95% posterior credible intervals, which indicate the range in which there is a .95 probability that each *r* value falls, as well as Bayes factors (BF_10_). BF_10_ is intuitively interpreted as an odds ratio for the research hypothesis; a value of 5, for example, indicates the data are 5 times more likely under the research hypothesis than under the null hypothesis. Although BF_10_ is best thought of as a continuous measure of evidence, values >3 are generally interpreted as substantial evidence for the research hypothesis and those <.33 as substantial evidence for the null (45). As we did not have an a priori expectation of whether correlations between model parameters and neural activations would be positive or negative, we simply tested the hypothesis that they were correlated in either direction (i.e., the *r* prior was set to be a uniform distribution between −1 and 1). We primarily used BF_10_ for inference due to its ability to quantify evidence for both the research and null hypotheses (45,46). However, we also report frequentist *p*-values, corrected for multiple comparisons with the False Discovery Rate (FDR=5%) method (47), to corroborate our BF_10_ inferences and assess whether correlations may have resulted from multiple testing.

Second, with the goal of indexing the latent constructs posited by the DDM at the neural level, we constructed latent variables for the neural correlates of any DDM parameters that were related to at least three ROIs. Latent variables were constructed by selecting the three ROIs that were most strongly linked to the parameter in the initial brain-behavior correlation analyses and entering them into a structural equation model in the R package *lavaan* (48). In these models, the three ROIs were treated as manifest measures of a neural latent variable that predicted the DDM parameter of interest in a regression. Factor scores from this latent variable were then extracted and were assumed to index the DDM construct at the neural level.

Third, with the goal of testing whether measures informed by DDM constructs predict psychopathology, we assessed correlational relationships between the behavioral and neural measures of these constructs and raw scores from two psychopathology scales on the ASR: externalizing behavior and DSM-IV attention-deficit/hyperactivity disorder (ADHD) symptoms. The former was selected because of previously-reported associations between go/no-go task-related neural activations and risk for substance use and other broad forms of externalizing behavior (14,23,28,29,49). The latter was selected because of the well-established associations between DDM parameters and ADHD diagnosis (33,50,51,52). For BF_10_, as we expected less efficient processing, less cautious response style, greater bias toward responding, and/or longer non-decision times to be related to increased psychopathology, we tested directional hypotheses by setting a uniform prior between 0 and 1 for indices we expected to be positively correlated with psychopathology (*z*, *Ter*) and a uniform prior between −1 and 0 for those we expected to be negatively correlated (*v*, *a*).

## Results

### fMRI Contrasts and ROI selection

Whole-brain maps for all four contrasts (Figure 2a) revealed neural responses in multiple regions that were broadly consistent with previous literature. The CR>GO contrast revealed right-lateralized activity in the frontal and parietal regions commonly inferred to be involved in top-down inhibitory control (18,53,54). The FA>GO and FA>CR contrasts identified areas associated with error processing and performance monitoring, including the ACC and bilateral clusters spanning the inferior frontal gyrus (IFG) and anterior insula (22,24).

**Figure 2.**
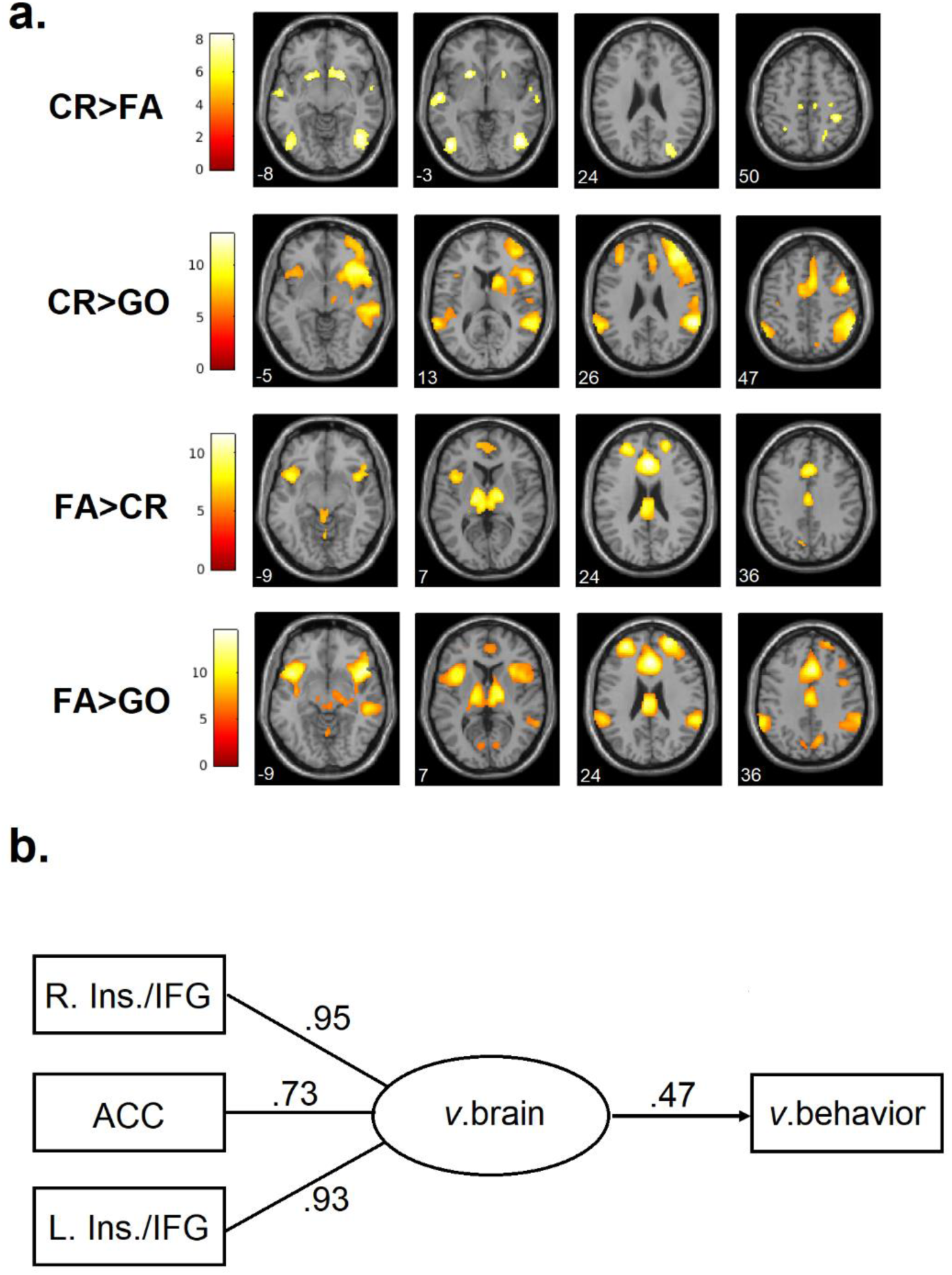
Neural activation in all fMRI contrasts and brain-based indices of the DDM parameter of drift rate (*v*). **a)** Whole-brain *t-*statistic maps (thresholded at FWE<.001) at selected axial slices for each contrast. White numbers in the bottom left corner indicate the z-coordinate for each slice. CR = correct rejection; FA = false alarm; GO = correct go trial. **b)** Model for the latent neural variable indexing drift rate (*v.*brain) with FA>CR activation estimates from the right anterior insula/inferior frontal gyrus (R. Ins./IFG), left anterior insula/inferior frontal gyrus (L. Ins./IFG), and anterior cingulate cortex (ACC). Standardized factor loadings and a standardized regression beta weight for prediction of average drift rate estimated with behavioral data (*v*.behavior) are displayed.

From these maps, we identified 23 frontal, parietal and sub-cortical ROIs (Table 2) that would be of the greatest interest given prior work (18,22,24,53,54) and extracted average parameter estimates for the respective contrasts from these regions. Similarity between the clusters identified by the FA>GO and FA>CR contrasts suggested that parameter estimates from ROIs in these contrasts were redundant; indeed, estimates from ROIs in the FA>GO contrast were highly correlated with those from the corresponding regions in the FA>CR contrast, ranging from *r*=.80 (right middle frontal gyrus) to *r*=.89 (anterior cingulate). Therefore, to reduce redundancy and limit the number of correlation tests, we only used ROIs drawn from the FA>CR contrast, for a final total of 16 ROIs in further analyses.

**Table 2.**
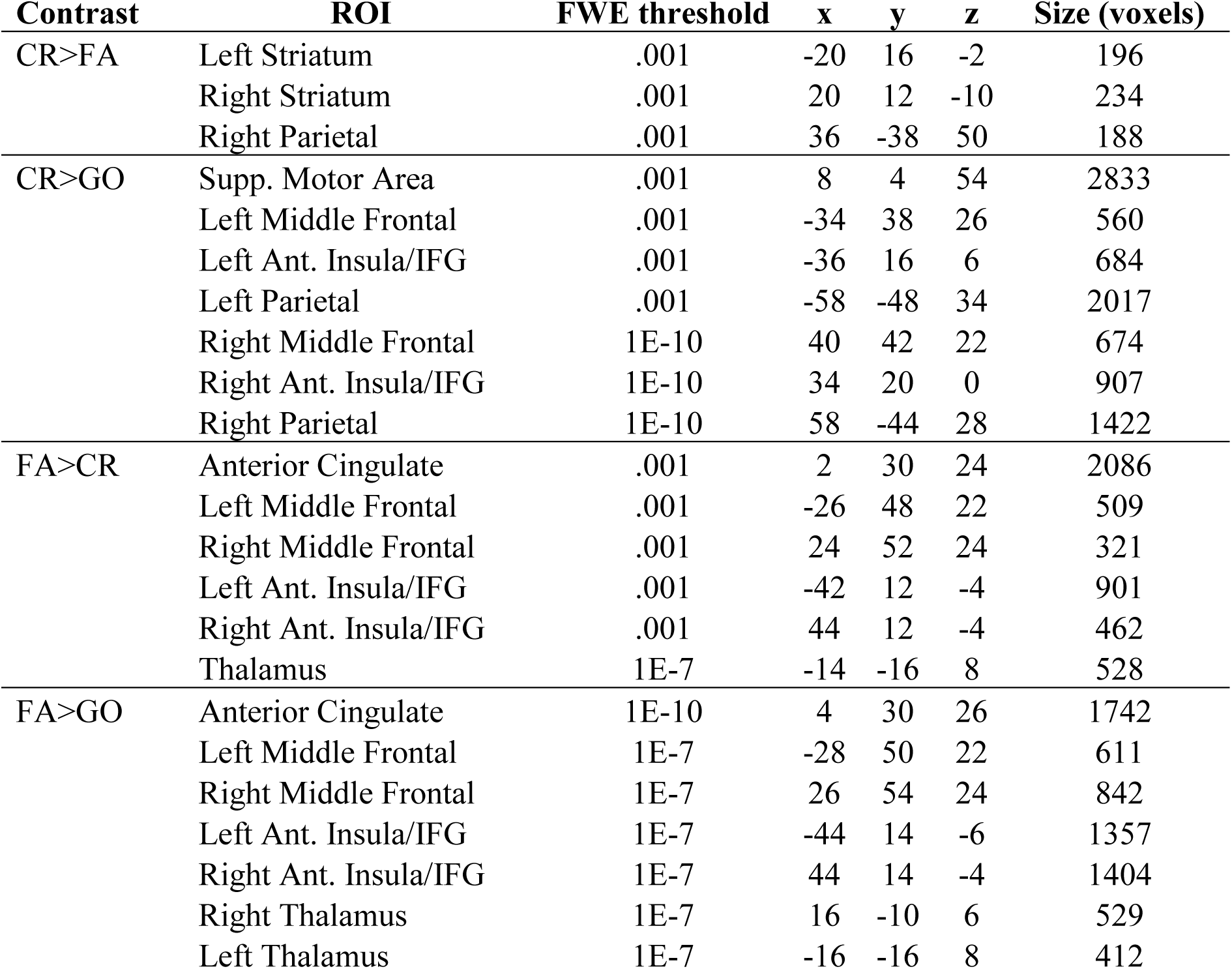
MNI coordinates, cluster-forming thresholds and size of all regions of interest (ROIs) selected from contrast maps for inclusion in further analyses. All ROIs were significant at a cluster-level threshold of FWE<.001, but stricter thresholds were used where noted to separate discrete clusters that were contiguous at this initial threshold. CR = correct rejection; FA = false alarm; GO = correct go trial; Ant. = Anterior; IFG = Inferior Frontal Gyrus; Supp. = Supplemental

| Contrast | ROI | FWE threshold | x | y | z | Size (voxels) |
| --- | --- | --- | --- | --- | --- | --- |
| CR>FA | Left Striatum | .001 | -20 | 16 | -2 | 196 |
|  | Right Striatum | .001 | 20 | 12 | -10 | 234 |
|  | Right Parietal | .001 | 36 | -38 | 50 | 188 |
| CR>GO | Supp. Motor Area | .001 | 8 | 4 | 54 | 2833 |
|  | Left Middle Frontal | .001 | -34 | 38 | 26 | 560 |
|  | Left Ant. Insula/IFG | .001 | -36 | 16 | 6 | 684 |
|  | Left Parietal | .001 | -58 | -48 | 34 | 2017 |
|  | Right Middle Frontal | 1E-10 | 40 | 42 | 22 | 674 |
|  | Right Ant. Insula/IFG | 1E-10 | 34 | 20 | 0 | 907 |
|  | Right Parietal | 1E-10 | 58 | -44 | 28 | 1422 |
| FA>CR | Anterior Cingulate | .001 | 2 | 30 | 24 | 2086 |
|  | Left Middle Frontal | .001 | -26 | 48 | 22 | 509 |
|  | Right Middle Frontal | .001 | 24 | 52 | 24 | 321 |
|  | Left Ant. Insula/IFG | .001 | -42 | 12 | -4 | 901 |
|  | Right Ant. Insula/IFG | .001 | 44 | 12 | -4 | 462 |
|  | Thalamus | 1E-7 | -14 | -16 | 8 | 528 |
| FA>GO | Anterior Cingulate | 1E-10 | 4 | 30 | 26 | 1742 |
|  | Left Middle Frontal | 1E-7 | -28 | 50 | 22 | 611 |
|  | Right Middle Frontal | 1E-7 | 26 | 54 | 24 | 842 |
|  | Left Ant. Insula/IFG | 1E-7 | -44 | 14 | -6 | 1357 |
|  | Right Ant. Insula/IFG | 1E-7 | 44 | 14 | -4 | 1404 |
|  | Right Thalamus | 1E-7 | 16 | -10 | 6 | 529 |
|  | Left Thalamus | 1E-7 | -16 | -16 | 8 | 412 |

### Neural Correlates of DDM Parameters

Plots comparing empirical RT and accuracy data to data predicted by the DDM suggested that the model generally described behavioral data well (Supplemental Materials). Hence, we determined that model fit was adequate, and proceeded to investigate links between DDM parameter estimates and neural activation. For these and all subsequent analyses, *v.go* and *v.nogo* were averaged to provide a general index of evidence accumulation efficiency (*v*). Notably, most participants’ start point (*z*) values were above .50 (Table 1), indicating that they were biased toward the decision to respond, relative to the decision to withhold a response, as would be expected for a task with a greater proportion of “go” relative to “no-go” stimuli.

Table 3 reports results from correlation tests of relationships between ROI activations and DDM parameters. The most substantial associations identified were the positive correlations between *v* and activity in the prefrontal regions identified by the FA>CR contrast. FA-related activations in regions putatively involved in error monitoring, the bilateral insula/IFG and ACC, were most strongly correlated with *v*. Neither efficiency of task processing (*v*) nor bias toward responding (*z*) were related to activity in the fronto-parietal network identified in the CR>GO contrast; Bayes factors generally suggested evidence *against* the presence of such relationships. Rather, activity in a subset of these ROIs displayed weak relationships with DDM parameters that index other processes; increased activity in the right middle frontal gyrus and right parietal lobe was related to a less cautious decision-making style (emphasizing speed over accuracy by lowering *a*), while increased activity in the latter was also linked to shorter non-decision times.

**Table 3.** Correlation (*r*) values, 95% posterior credible intervals (CIs), Bayes factors (*BF_10_*) and frequentist *p*-values for correlational relationships between ROI activations and DDM parameters. For Bayes factors: **bolded** = substantial evidence (>3:1 odds) for a correlational relationship; For *p*-values: *=survives FDR correction for multiple comparisons within each family of tests (families defined by DDM parameters)

| Region | Efficiency ($v$) | | | Bias ($z$) | | |
| --- | --- | --- | --- | --- | --- | --- |
| | $r$ [CI] | $BF_{10}$ | $p$ | $r$ [CI] | $BF_{10}$ | $p$ |
| CR/FA Left Striatum (-20,16,-2) | -.10[-.25,.07] | 0.2 | .256 | .09[-.08,.25] | 0.18 | .305 |
| CR/FA Right Parietal (36,-38,50) | -.16[-.32,.00] | 0.66 | .053 | -.10[-.26,.06] | 0.22 | .219 |
| CR/FA Right Striatum (20,12,-10) | -.11[-.26,.06] | 0.23 | .208 | .11[-.06,.27] | 0.24 | .191 |
| CR/GO Left Ant. Insula/IFG (-36,16,6) | -.14[-.29,.03] | 0.37 | .109 | .05[-.11,.21] | 0.12 | .557 |
| CR/GO Left Middle Frontal (-34,38,26) | .05[-.12,.21] | 0.12 | .584 | .04[-.13,.20] | 0.11 | .679 |
| CR/GO Left Parietal (-58,-48,34) | -.18[-.33,-.01] | 0.9 | .037 | .12[-.05,.27] | 0.27 | .165 |
| CR/GO Right Ant. Insula/IFG (34,20,0) | .01[-.15,.17] | 0.11 | .901 | .11[-.06,.27] | 0.25 | .190 |
| CR/GO Right Middle Frontal (40,42,22) | .13[-.03,.29] | 0.36 | .114 | .10[-.06,.26] | 0.22 | .217 |
| CR/GO Right Parietal (58,-44,28) | .09[-.08,.24] | 0.17 | .314 | .07[-.10,.23] | 0.14 | .430 |
| CR/GO Supp. Motor Area (8,4,54) | .03[-.13,.19] | 0.11 | .713 | .03[-.14,.19] | 0.11 | .747 |
| FA/CR Anterior Cingulate (2,30,24) | .40[.25,.53] | **2.1E+4** | 6.7E-7* | -.11[-.27,.05] | 0.26 | .178 |
| FA/CR Left Ant. Insula/IFG (-42,12,-4) | .41[.26,.53] | **3.9E+4** | 3.5E-7* | -.11[-.27,.05] | 0.26 | .179 |
| FA/CR Left Middle Frontal (-26,48,22) | .27[.11,.41] | **20.71** | 1.1E-3* | -.04[-.20,.12] | 0.12 | .631 |
| FA/CR Right Ant. Insula/IFG (44,12,-4) | .45[.30,.57] | **6.4E+5** | 1.9E-8* | -.15[-.30,.01] | 0.52 | .073 |
| FA/CR Right Middle Frontal (24,52,24) | .36[.20,.49] | **1.4E+3** | 1.2E-5* | -.15[-.31,.01] | 0.55 | .067 |
| FA/CR Thalamus_(-14,-16,8) | -.02[-.18,.15] | 0.11 | .852 | -.20[-.35,-.04] | 1.89 | .016 |

| Region | Caution ($a$) | | | Non-decision ($Ter$) | | |
| --- | --- | --- | --- | --- | --- | --- |
| | $r$ [CI] | $BF_{10}$ | $p$ | $r$ [CI] | $BF_{10}$ | $p$ |
| CR/FA Left Striatum (-20,16,-2) | -.18[-.33,-.02] | 1.14 | .028 | -.24[-.38,-.08] | **6.10** | .004* |
| CR/FA Right Parietal (36,-38,50) | -.04[-.20,.12] | 0.12 | .634 | .00[-.16,.16] | 0.11 | .988 |
| CR/FA Right Striatum (20,12,-10) | -.21[-.36,-.05] | 2.31 | .012 | -.25[-.39,-.09] | **8.50** | .003* |
| CR/GO Left Ant. Insula/IFG (-36,16,6) | -.16[-.31,.01] | 0.62 | .058 | -.09[-.25,.07] | 0.19 | .267 |
| CR/GO Left Middle Frontal (-34,38,26) | -.20[-.35,-.04] | 1.79 | .017 | -.03[-.19,.13] | 0.11 | .711 |
| CR/GO Left Parietal (-58,-48,34) | -.20[-.35,-.03] | 1.54 | .020 | -.17[-.32,.00] | 0.73 | .047 |
| CR/GO Right Ant. Insula/IFG (34,20,0) | -.19[-.34,-.03] | 1.43 | .022 | -.16[-.32,.00] | 0.67 | .053 |
| CR/GO Right Middle Frontal (40,42,22) | -.23[-.37,-.06] | **3.84** | .007 | -.05[-.21,.12] | 0.12 | .570 |
| CR/GO Right Parietal (58,-44,28) | -.24[-.39,-.08] | **6.52** | .004 | -.22[-.37,-.06] | **3.27** | .008* |
| CR/GO Supp. Motor Area (8,4,54) | -.15[-.31,.01] | 0.54 | .069 | -.07[-.23,.10] | 0.14 | .423 |
| FA/CR Anterior Cingulate (2,30,24) | .04[-.13,.20] | 0.12 | .659 | .09[-.08,.24] | 0.17 | .312 |
| FA/CR Left Ant. Insula/IFG (-42,12,-4) | .05[-.12,.21] | 0.12 | .586 | .16[.00,.31] | 0.64 | .055 |
| FA/CR Left Middle Frontal (-26,48,22) | -.06[-.22,.10] | 0.14 | .448 | .07[-.09,.23] | 0.15 | .389 |
| FA/CR Right Ant. Insula/IFG (44,12,-4) | .11[-.06,.27] | 0.24 | .197 | .19[.03,.34] | 1.31 | .024 |
| FA/CR Right Middle Frontal (24,52,24) | .01[-.15,.17] | 0.11 | .921 | .06[-.10,.22] | 0.14 | .453 |
| FA/CR Thalamus_(-14,-16,8) | -.12[-.28,.04] | 0.29 | .150 | .11[-.05,.27] | 0.26 | .180 |

### Brain and Behavioral Measures of DDM Constructs

As the strongest links between DDM parameters and ROI activation were between *v* and FA-related activation in the putative error monitoring network involving the ACC and bilateral insula/IFG, we constructed a structural equation model (Figure 2b) in which these three regions were manifest indices of a latent neural variable (*v*.brain) that predicted DDM estimates of *v* from behavioral data (*v*.behavior). Practical fit indices suggested that the model described the data adequately (CFI=.999, TLI=.998, RMSEA=.031, SRMR=.020), and all relationships in the model, including the three factors loadings and the regression in which *v*.brain predicted *v*.behavior, displayed *p-*value*s* <.001. Although the *Ter* parameter was weakly linked to activity in three ROIs, a similar model in which activity in these ROIs formed a factor to predict *Ter* displayed questionable fit (CFI= .98, TLI=.95, RMSEA=.102, SRMR=.047) and abnormalities (negative variance estimates) that suggested it was poorly specified. Therefore, the primary measures of interest were indices of the construct of evidence accumulation efficiency at the behavioral (*v*.behavior) and neural (*v*.brain) levels. To ensure that our ROI selection strategy did not bias predictions of psychopathology, we also conducted a sensitivity analysis (Supplemental Materials) where we used the first principle component of neural responses in the FA>CR contrast as an alternate *v*.brain measure. This component was strongly correlated with the *v*.brain measure reported here (*r*=.92) and displayed similar relationships with *v*.behavior and psychopathology.

### Prediction of Psychopathology

Correlation tests of relationships between model-based measures and clinical outcomes of interest (externalizing behaviors, ADHD symptoms) are displayed in Table 4. Although *v*.behavior did not predict self-reported externalizing behavior, there was moderate evidence that this index was negatively related to ADHD symptoms, consistent with prior case-control studies. However, *v*.brain displayed moderate to strong evidence of negative relationships with both outcomes. All other DDM parameters displayed evidence *against* the presence of hypothesized relationships with psychopathology.

**Table 4.** Correlation (*r*) values, 95% posterior credible intervals (CIs), Bayes factors (*BF_10_*) and frequentist *p*-values for relationships between model-based indices and externalizing psychopathology self-ratings. All analyses excluded the 5 subjects who did not have ASR data available. CIs and *r* values were estimated assuming a uniform prior from −1 to 1. However, as we specifically wished to test the hypotheses that less efficient neurocognitive functioning and/or less cautious response style were related to increased rates of psychopathology, Bayes factors for *v*.behavior, *v*.brain, and *a* tested the directional hypothesis that the relationship with psychopathology was negative (i.e., the uniform prior spanned values between −1 and 0). As we wished to test the hypotheses that higher levels of bias toward responding and/or longer non-decision times were related to increased rates of psychopathology, Bayes factors for *z* and *Ter* tested the hypothesis that the relationship with psychopathology was positive (i.e., the uniform prior spanned values between 0 and 1). For Bayes factors: **bolded** = substantial evidence (>3:1 odds) for the tested correlational relationship; For *p*-values: *=survives FDR correction for multiple comparisons within each family of tests (families defined by psychopathology measure)

| Predictor | Externalizing Scale (raw) |  |  | DSM-ADHD Scale (raw) |  |  |
| --- | --- | --- | --- | --- | --- | --- |
| | $r$ [CI] | $BF_{10}$ | $p$ | $r$ [CI] | $BF_{10}$ | $p$ |
| $v$.behavior | -.13[-.29,.04] | .64 | .124 | -.21[-.36,-.05] | **4.70** | .012* |
| $v$.brain | -.23[-.38,-.06] | **7.90** | .007* | -.28[-.42,-.12] | **48.29** | 9.3E-04* |
| $a$ | -.07[-.24,.09] | .25 | .393 | -.07[-.23,.10] | .23 | .418 |
| $z$ | -.06[-.22,.11] | .07 | .497 | -.03[-.19,.14] | .08 | .749 |
| $Ter$ | -.03[-.20,.14] | .08 | .716 | .04[-.12,.21] | .17 | .614 |

## Discussion

The current study assessed whether activations from several common go/no-go neuroimaging contrasts were related to latent cognitive processes indexed by the DDM, a well-validated mathematical model of the go/no-go task (30,31), and whether these activations and corresponding cognitive processes were related to externalizing psychopathology. This approach was aimed at testing the common assumption in the clinical neuroscience literature that individual differences in go/no-go task-related neural activations index the integrity of clinically-relevant neurocognitive mechanisms.

Individual differences in CR-related fronto-parietal activations, which are generally described as the neural substrate of response inhibition (18,53,54), were neither related to individuals’ efficiency at deciding whether or not to withhold a response (*v*) nor related to their level of bias toward responding relative to withholding (*z*). Instead, greater activation in the right middle frontal gyrus and right parietal lobe was weakly related to less cautious decision-making styles (lower *a*). Although lower *a* may be interpreted as more “impulsive” responding, and increased activation during CRs could therefore be posited as a neural marker of impulsivity, this interpretation is tenuous for two reasons. First, the relationships observed between *a* and activations were relatively weak for relationships between variables that presumably index the same construct, and were not robust to corrections for multiple testing. Second, as *a* did not predict externalizing behaviors or ADHD symptoms, individual differences in caution on the standard go/no-go task appear to be unrelated to these clinical entities, despite the fact that these entities are generally associated with impulsivity. Therefore, the current findings cast doubt on the conventional interpretation of CR-related fronto-parietal activations as corresponding to individual differences in inhibition ability, and suggest that if they do index individual differences in a mechanistic process, such as caution, that process is likely to be orthogonal to externalizing behavior.

In contrast, FA-related activations in the ACC and bilateral insula, regions related to error processing (22,24), were strongly correlated with *v*, suggesting that neural systems involved in performance monitoring are crucial for optimizing an individual’s ability to efficiently decide whether to initiate or withhold responses. These activations also displayed evidence of relationships with clinical outcomes; the neural correlates of *v* predicted individual differences in both externalizing behaviors and ADHD symptoms. Taken together with the body of behavioral research linking ADHD and externalizing behaviors to poorer performance on the go/no-go task (1,2,55,56), these results suggest that the task does indeed tap into a neurocognitive construct with clinical relevance. However, rather than being exclusively related to the inhibition of responses or to neural processes that are engaged on CR trials only, the current findings suggest that this construct may be better-defined as a general efficiency of evidence accumulation, which may be dependent on performance monitoring systems.

Such an explanation is consistent with recent accounts of cognitive deficits in ADHD. Although response inhibition has long been considered a core deficit in the disorder (57), individuals with ADHD display lower rates of evidence accumulation than their peers across a wide variety of tasks that vary in their response inhibition demands (52,58,59). These findings have recently been interpreted by some (58,59) as suggesting that individuals with ADHD display dysfunction related to the locus coeruleus norepinephrine (LC-NE) system, which is posited (60) to optimize arousal and the efficiency of processing in response to perceived task utility. This account implies that lower evidence accumulation efficiency may reflect either dysfunction in the LC-NE system itself, dysfunction in top-down inputs to the system that modulate its activity in response to information about task utility and performance, or lower perceived task utility due to broader factors. As the ACC and other brain regions involved in monitoring performance lapses are thought to provide top-down input to the LC-NE system to enhance task processing (60), the current study’s findings on the neural correlates of *v* are highly consistent with this account.

The current study has several limitations. First, the study only assessed correlational relationships between behavioral indices, neural activations, and clinical scales from the same time point. Future work involving prospective prediction of clinical outcomes would be useful for clarifying whether model-based measures from the go/no-go task provide information about individuals’ predisposition to psychopathology. Second, as the majority of the sample was male and many individuals displayed existing risk factors for externalizing psychopathology, our findings may not generalize to samples without these features. Third, although the DDM parameters and their associated neural activations predicted clinical outcomes, the effect sizes were small, which has implications for the clinical utility of this work. Finally, an inherent limitation of any work involving a formal cognitive model is that the conclusions drawn may be peculiar to the assumptions of the specific model used. For example, if another model was used which explained response inhibition on “no-go” trials with a set of mechanisms separate from those assumed by the DDM, it is possible that different conclusions about the neural correlates of response inhibition would have been reached. However, as the DDM provides a comprehensive description of task performance on both “go” and “no-go” trials with a parsimonious set of parameters, it is reasonable assume that the basic DDM adequately explains performance on the task without invoking separate “inhibition” mechanisms.

In conclusion, the current study assessed relationships between neural activations from common go/no-go task contrasts and parameter estimates from the DDM to test the assumption that neural responses in these contrasts index individual differences in the integrity of clinically-relevant neurocognitive processes. Surprisingly, activation in the right-lateralized fronto-parietal network associated with successful inhibition was not related to the integrity of cognitive processing. In contrast, activity during inhibitory errors in the ACC and bilateral insula was strongly related to efficiency of processing on the task and predicted externalizing behaviors and ADHD symptoms. These results call common mechanistic interpretations of go/no-go task-related activations into question and suggest that these activations can inform clinical neuroscience by providing information about neural systems that monitor and optimize task performance in response to perceived task utility, rather than about the specific neural correlates of withholding responses.

## Acknowledgements

This project was supported by NIAAA grants R01 AA07065 and R01 AA025790. Alexander Weigard was supported by NIAAA T32 AA007477.

## Supplemental Materials

### Neuroimaging Data Acquisition Parameters

Whole brain T2*-weighted MRI images were acquired on a 3.0 T GE Signa scanner (Milwaukee, WI) using a single-shot spiral in-out sequence (1) with the following parameters: TR=2000ms, TE=30ms, flip angle=90°, FOV=200mm, 29 axial slices, 64×64 matrix, in-plane resolution=3.12mm×3.12mm, and slice thickness=4mm. To assist with inter-subject spatial normalization, a high-resolution T1-weighted anatomical image was acquired in a separate scan with the following parameters: three-dimensional spoiled gradient-recalled echo, TR=25ms, minimum TE, FOV=25cm, 256×256 matrix, slice thickness=1.4mm.

### Diffusion Decision Model Specification and Parameter Estimation

The current study focused on MLS participants who completed their first session of the neuroimaging study when they were ages 18-21. However, the neuroimaging component of the MLS has recruited additional participants since these initial sessions (including the offspring of the original MLS participants), and has attempted to conduct longitudinal neuroimaging data collection at 1- to 2-year intervals with as many individuals as possible. Therefore, at the time that analyses for the current project began, data from a total of 1280 go/no-go neuroimaging sessions had been collected from 306 individual participants while they were between the ages of 7 and 30. Rather than only fitting the diffusion decision model (DDM) to data from the neuroimaging sessions used for the current study, we decided to fit the DDM to all sessions from the MLS neuroimaging sample with valid go/no-go behavioral data for two reasons. First, doing so provided us with many more data points with which to evaluate model fit and to enter into our simulation-recovery study to assess the reliability of parameter estimates. Second, we aimed to produce parameter estimates that could be leveraged in future work involving the other time points from the neuroimaging study. Of the 1280 sessions available, 1255 met our inclusion criteria for data quality, which were 1) that at least 200 trials were available for model-fitting and 2) that the overall accuracy rate in the session was greater than .55, indicating that the participant understood, and was engaging in, the task. After selection of the included sessions, response times (RTs) less than 200ms were excluded from analysis as fast guesses, following standard procedures for fitting the DDM (2), and the DDM was fit to each session separately.

As noted in the main text, “simple” versions DDM, which do not include the parameters for between-trial variability in drift rate (*sv*), start point (*sz*) or non-decision time (*st0*), are often preferable to “full” versions of the DDM for two reasons. First, as it is difficult to reliably estimate these between-trial variability parameters without very large numbers of trials, fixing them to 0 for most applications of the DDM likely makes estimates of the main model parameters (*v*, *a*, *z*, *Ter*) more stable (3). Second, evidence from a blinded, collaborative test of researchers’ ability to draw valid inferences from response time models (4) strongly suggested that simple versions of the DDM provided inferences about constructs indexed by the main DDM parameters that were just as robust and valid those provided by full versions of the DDM. Therefore, we first fit a “simple” version of the DDM which only contained 5 parameters: drift rate for “go” stimuli (*v.go*), drift rate for “no-go” stimuli (*v.nogo*), boundary separation (*a*), non-decision time (*Ter*). Decisions to respond on a given trial were assumed to occur when the diffusion process crossed the upper response boundary, while decisions to withhold from responding were assumed to occur when the process crossed the lower response boundary, which was equivalent to the “implicit” boundary assumed by (5,6). Hence, more positive values of *v.go* indicate more efficient accumulation of evidence for the correct response on “go” trials, but more negative values of *v.nogo* indicate more efficient accumulation of evidence for the correct response on “no-go” trials. For all subsequent analyses, *v.nogo* parameter estimates were multiplied by −1 so that they would be directly comparable with *v.go* parameter estimates. The *z* parameter was parameterized as a proportion of *a*, meaning that values above .5 indicate a bias toward responding and values below .5 indicate a bias to non-responses.

The model was in fit by implementing the chi-square minimization procedure specified for the go/no-go DDM by (5) and (6) using functions from the R package *rtdists* (7) and base functions in the R language. Chi-square values to be minimized were calculated for each set of model parameters by first calculating the proportion of responses that the model predicted would terminate at the upper and lower response boundaries, as well as predicted RT quantiles (.1, .2, .3, .4, .5, .6, .7, .8, and .9) for correct responses to “go” stimuli and erroneous responses to “no-go” stimuli, which formed 10 bins for response times (e.g., a bin for responses less than the .1 quantile, a bin for responses between the .1 and .2 quantiles, etc.). The expected proportion of response times in each bin was .1 multiplied by overall proportion of each response type (respond/withhold) predicted by the model for each condition (“go”/”no-go”), which produces expected proportions that are weighted by accuracy in each condition. As non-responses in the “go” and “no-go” conditions do not have observed response times, a single bin was used. The expected proportion of responses in this bin was simply 1 multiplied by the proportion of non-responses predicted by the model in each condition. Following prior work (Ratcliff et al., 2018), we used an alternate RT binning procedure when the number of RTs in a given condition was small: when the number of RTs was <11 we used the predicted median RT to form two bins (where the expected proportion was .5 multiplied by the expected proportion for that response), and when the number of trials was <4, we used a single bin (the same procedure as was used for non-responses). The expected number (E) of responses for each of the bins was then calculated by multiplying the response and RT proportions predicted by the model by the actual number of trials in each stimulus condition. The observed number (O) of responses in each bin was then counted, and a chi-square value for each bin was calculated as (O - E)^2^/E. Overall chi-square values were then calculated by summing over all bins for the individual.

With the overall chi-square value as an optimization criterion, the R function optim() was used to implement a multidimensional search using the Nelder-Mead method. Following previous work (8), starting points for the initial search process were found using the EZ diffusion model (9) to estimate *v*, *a*, and *Ter* parameters for data in the “go” condition. The starting value for *v.go* was the *v* estimate from EZ and the starting value for *v.nogo* was this estimate multiplied by −1. Start points for the *a* and *Ter* parameters were equivalent to the respective parameters from the EZ fits, and the start point for *z* was always set at .5, as EZ does not estimate response bias. Running the Nelder-Mead algorithm multiple times and using parameter estimates from each run as start points for the following run often notably improves model fit (8). Therefore, this procedure was adopted for the current analysis; optim() was set to run as many times as needed until no further decrease in the chi-square value could be accomplished (mean number of minimization runs per neuroimaging session = 5.07, SD = 2.35). Bounds were placed in the search space for several parameters to prevent impossible values (*z*>1,*z*<0, *a*<0,*Ter*<0) or unrealistically large values (*v* > 6, *v* < −6) by setting the objective function to return an infinite chi-square if such a parameter value is entered.

### Assessment of Model Fit

Model fit was assessed by plotting model-predicted accuracy rates and RT quantiles (.1, .5, .9) for the “go” and “no-go” stimulus conditions against actual values of the same accuracy rates and RT quantiles for each go/no-go task session. Supplemental Figures 1 and 2 display these plots for data from every neuroimaging session in the MLS sample with useable data (N=1255), and separately for the 143 sessions that were further analyzed in the current study. In these plots, points clustered around the diagonal indicate good model fit. Points farther from the diagonal represent misfits, and cases in which the majority of points fall above or below the diagonal indicate a bias, as they suggest that the model is either over- or under-predicting the RT or accuracy rates in a given condition. Inspection of the plots indicates that the model provided an excellent description of performance on “go” trials; most points are clustered close to the diagonal for both RT quantiles and accuracy rates. For “no-go” trials, although points generally clustered around the diagonal, there was relatively more misfit, and an apparent bias in which the model systematically over-predicted RTs for false alarms, which was most pronounced for the longest RTs (.9 quantile). This misfit likely reflects the challenge of describing RT data in this condition, which are much sparser than RT data in the “go” condition. The vast majority of subjects had accuracy rates greater than .50 on “no-go” trials, indicating that less than 30 RTs were available for fitting in this condition, which may explain why model predictions for some “no-go” RT quantiles are less accurate. Nonetheless, predicted “no-go” RT quantiles were still highly correlated with empirical quantiles (*r* = .90, .89 and .72 for the .1, .5 and .9 “no-go” RT quantiles, respectively, in the full N=1255 sample), suggesting that the model described individual differences in “no-go” RT data relatively well under the circumstances. Therefore, we concluded that model fit was adequate.

### Simulation-Recovery Study to Assess DDM Parameter Reliability

To assess whether the model and fitting method used could be expected to reliably recover DDM parameters given the number of trials in the MLS go/no-go task, we conducted a simulation/recovery study. First, 400 of the 1255 task sessions from the MLS sample that were fit to the DDM were randomly selected. Next, parameters from each of these sessions were used to simulate 400 separate data sets which each consisted of the same number of “go” stimulus trials (n=185) and “no-go” stimulus trials (n=60) as the empirical data. The DDM was then fit to all 400 simulated data sets using the same procedures outlined above and parameter estimates recovered from the data sets were compared with those used to simulate the data. We adopted the convention for determining quality of recovery that was used by (10); correlation (*r*) values for the relationship between the simulated and recovered parameters were considered “poor/unacceptable” if *r* < .50, “fair” if .50 < *r* < .75, “good” of .75 < *r* < .90 and “excellent” if *r* > .90. Supplemental Figure 3 displays scatterplots and *r* values for all parameters, which indicate that every parameter displayed “good” or “excellent” recovery except in the case of *z*, where recovery was “fair”. Hence, this analysis provided evidence that the DDM displayed acceptable parameter recovery when fit to the data in this sample using the procedures outlined above.

### Correlations Between ROI Activations and Behavioral Summary Statistics

Of the behavioral summary statistics commonly used to index task performance in prior work (Supplemental Table 1), FA rate and hit SDRT were strongly linked to the putative error-processing-related activations in the FA>CR contrast that were also linked to *v*. However, these measures also showed associations with a handful of regions in the CR contrasts, with worse performance (higher FA rates) generally being linked to greater activation. Increased CR>GO contrast activation in right fronto-parietal regions was also related to faster RTs, although the associations, reported in the main body of the manuscript, of the same regions with the *a* parameter of the DDM suggest that this relationship with MRT is due to individual differences in response caution (less cautious responding with greater activation) rather than individual differences in the integrity of task performance.

### Sensitivity Analysis with Alternate Measure of *v* at the Neural Level

Our primary strategy for obtaining measures of DDM parameters at the neural level, as described in the text, involved 1) entering the three ROIs that were most strongly related to the parameter into a structural equation model in which these ROIs created a latent factor that predicted the parameter, and 2) extracting factor scores as the neural-level measure (i.e., *v*.brain). Although the procedure was specifically designed to identify patterns of neural activity that were most closely related to the individual DDM parameters, it could be argued that this approach would bias the neural measure’s prediction of psychopathology; if parameter estimates drawn from behavioral data are related to psychopathology, then ROIs that are selected precisely because they are related to these parameter estimates may show correlations with psychopathology that are artificially inflated. In order to address this concern, we conducted a sensitivity analysis that involved a more data-driven approach to identifying neural measures of *v*, the only parameter that appeared to be robustly related to brain responses.

First, given that *v* was exclusively related to neural responses in the FA>CR contrast, we conducted a principle component analysis (PCA) of the activation estimates of all ROIs from this contrast using the R package *FactoMineR* (11). Results of this PCA are displayed in Supplemental Table 2a. The first component, which explained over 63% of the variance, was most strongly correlated with the putative error monitoring regions that were entered in to the *v*.brain latent variable model in the main text: the ACC and bilateral insula. The first component was correlated with *v*.behavior (*r*=.41, *p*<.001) and very strongly correlated with the *v*.brain latent variable obtained using our original procedure in the main text (*r*=.92, *p*<.001), suggesting that it reflects an individual difference in neural function that can similarly be thought of a neural-level measure of *v*. Next, we assessed relationships of this measure with the clinical outcomes of interest. Similar to correlation tests involving the *v*.brain latent variable from the main text, we used Bayes factors (BF_10_) to test the directional hypothesis that the first FA>CR component was negatively related to psychopathology. We also corrected for multiple comparisons (including correlation tests involving *v*.behavior, the original *v*.brain, and the other DDM parameters) with the False Discovery Rate (FDR=5%) method. Results (Supplemental Table 2b) indicate moderate evidence that the FA>CR component is negatively related to both externalizing behavior and ADHD symptoms, and that the strength of these negative correlations is similar to that of the correlations between *v*.brain and the same clinical outcomes.

Taken together, results of this sensitivity analysis suggest that the component that explains the majority of the variance in the FA>CR contrast is highly similar to the *v*.brain measure obtained via the ROI selection procedure used in the main text, and can be similarly thought of as a neural-level measure of *v*. As there was evidence that this measure was also similarly related to the clinical outcomes of interest, we concluded that the ROI selection procedure used in the main text to create a neural-level index of *v* did not produce spurious relationships with psychopathology.

**Supplemental Figure 1.**
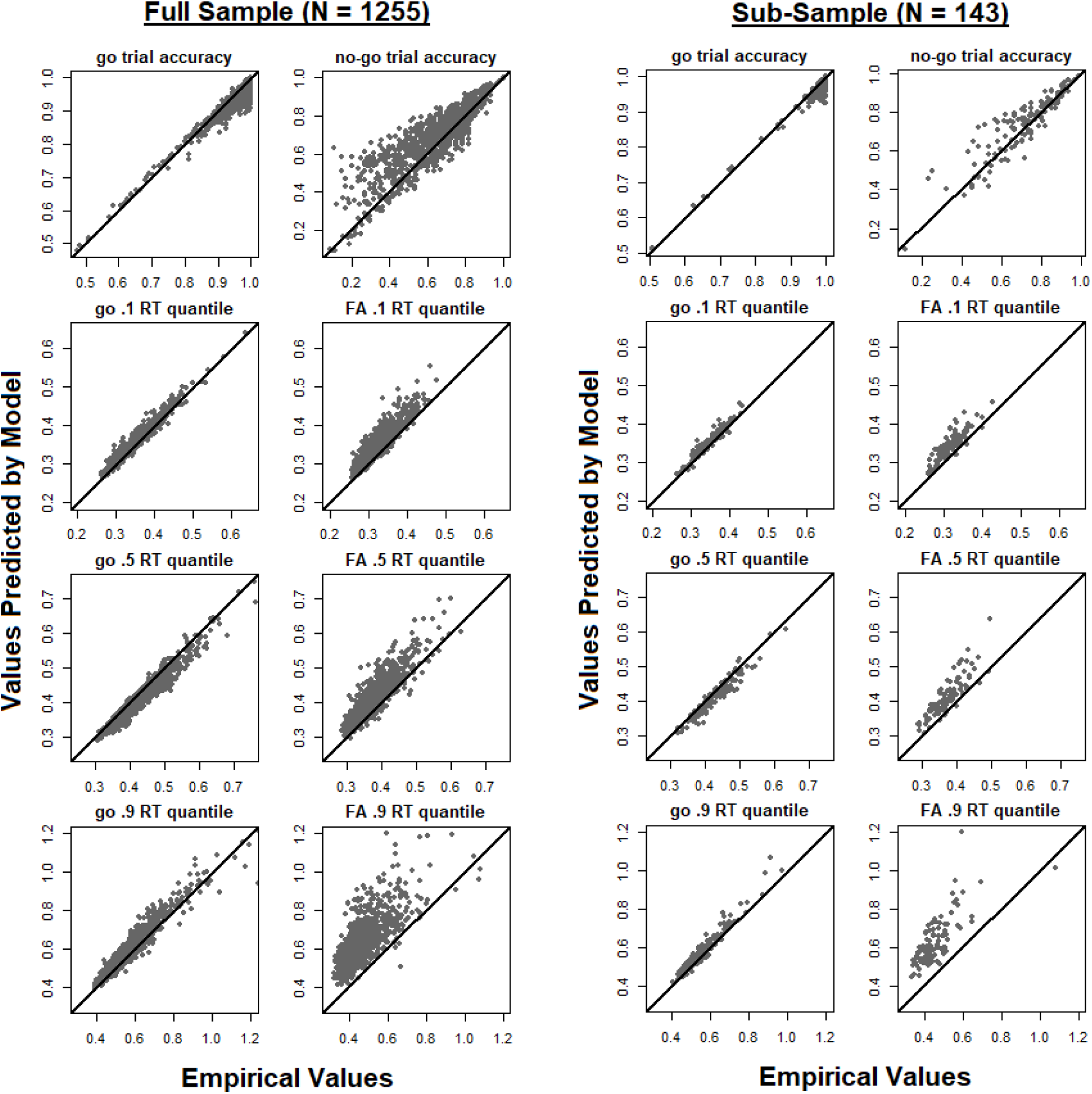
Empirical data for “go” and “no-go” condition accuracy rates and correct go and false alarm (FA) RT quantiles (.1, .5 and .9) plotted against the same values predicted by the model. The left panel displays data from all 1255 go/no-go task sessions entered into the analysis while the right panel only displays data for the 143 sessions that were analyzed further in the current study. Following previous work (Ratcliff, Huang-Pollock & McKoon, 2018) FA RT quantile data are only displayed for sessions with >10 observed RTs. The diagonal line indicates where points would fall if there was a perfect relationship between the empirical and predicted values. For clarity and comparability between the larger sample and sub-sample, axis intervals are set to be the same between samples and between “go” and “no-go” RT quantiles. Therefore, these plots do not include several outlier RT quantile values, which are shown in plots displayed in Supplemental Figure 2, below.

**Supplemental Figure 2.**
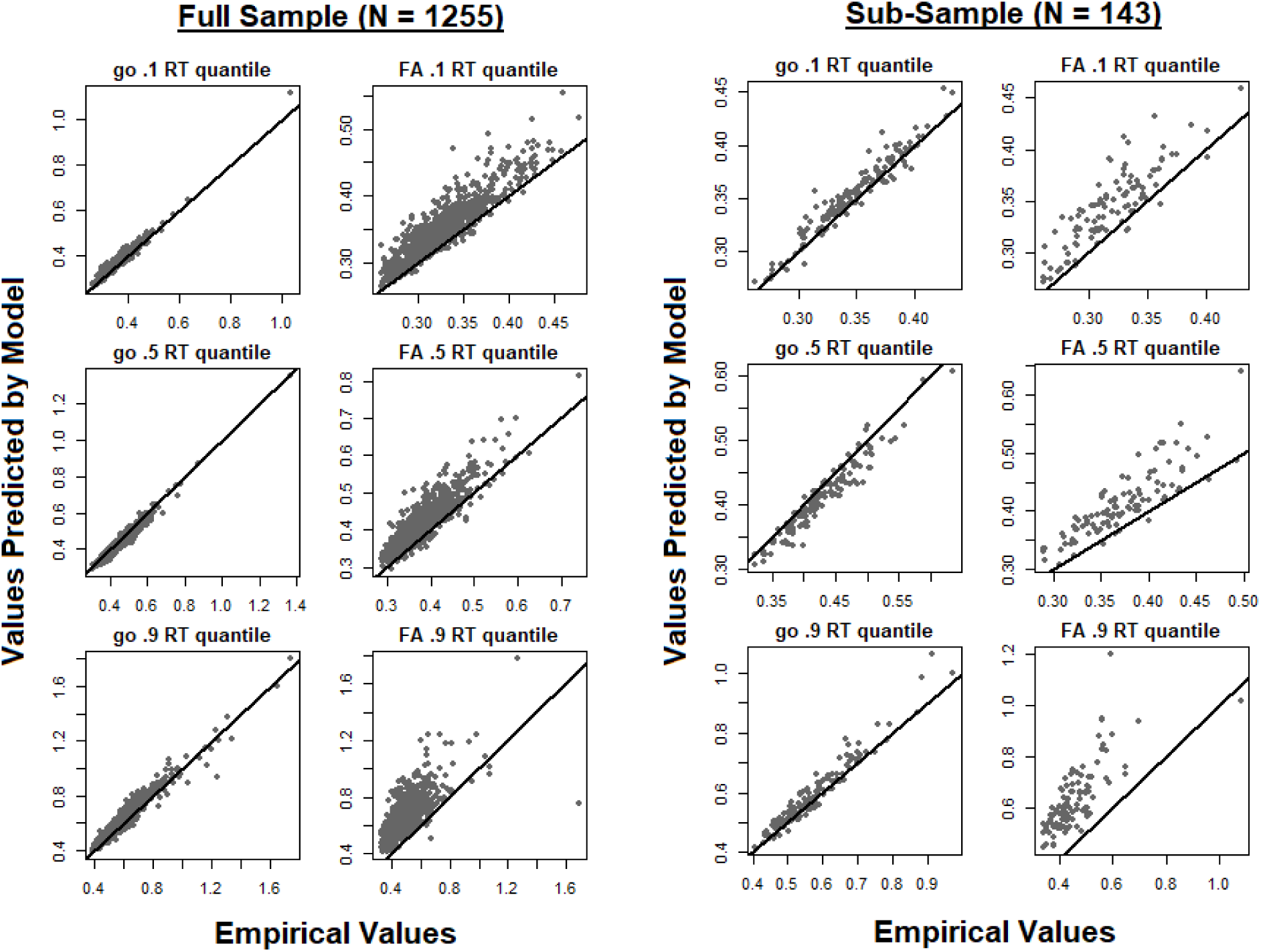
The same RT quantile data as displayed in Supplemental Figure 2 with plot axes adjusted to accommodate all data in each specific condition. As in in the previous figure, the diagonal line indicates where points would fall if there was a perfect relationship between the empirical and predicted values.

**Supplemental Figure 3.**
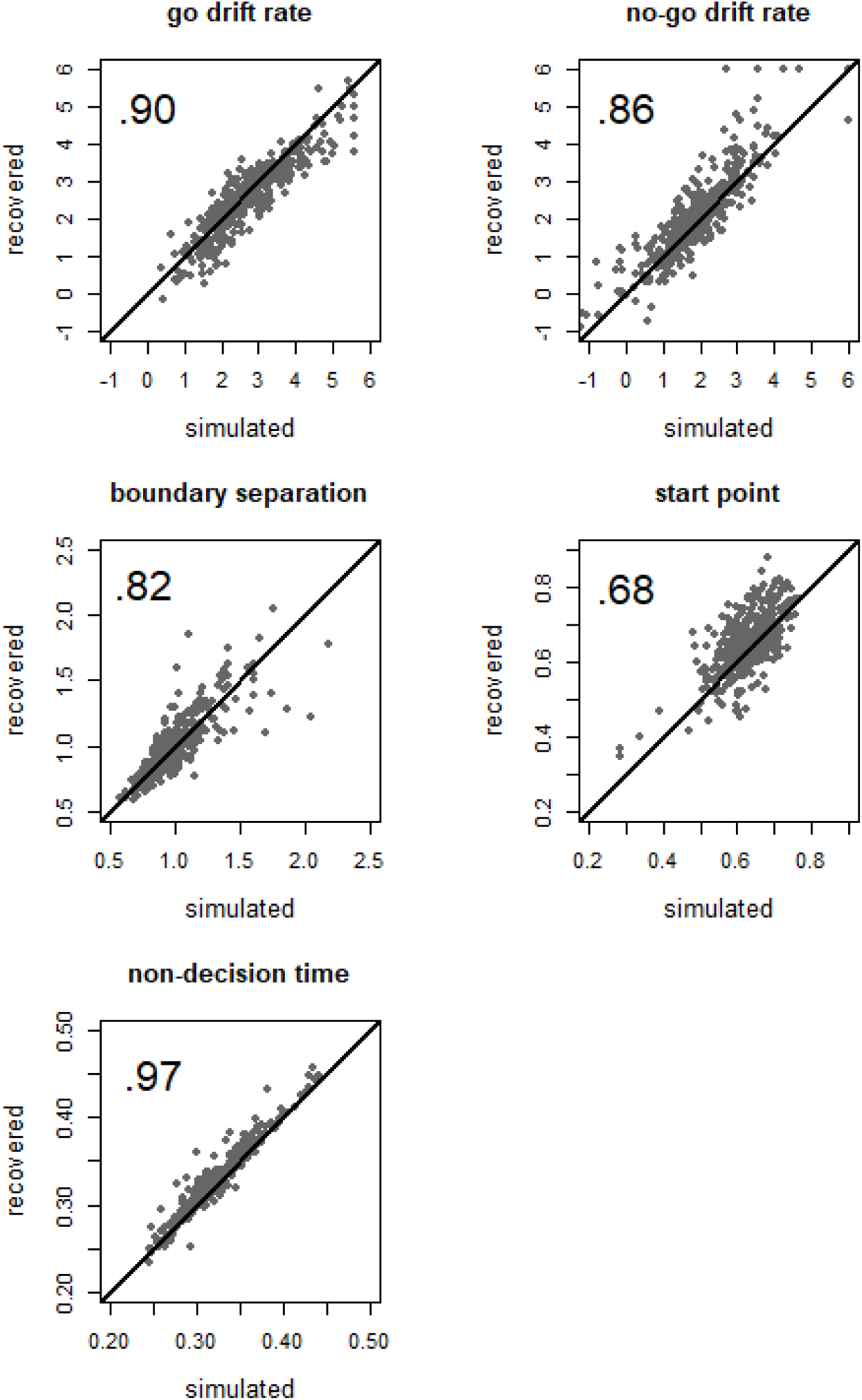
Parameter values used to simulate 400 data sets (drawn at random from 400 of the 1255 actual task sessions) plotted against parameter values that were recovered when these data sets were fit the DDM using the procedures outlined above. Correlation values (*r*) between the simulated and recovered parameter values are displayed in the top left corner of each plot Diagonal lines indicate where points would fall if there was perfect parameter recovery.

**Supplemental Table 1.** Correlation (*r*) values, 95% credible intervals (CIs), Bayes factors (*BF_10_*) and frequentist *p*-values for correlational relationships between ROI activations and behavioral summary statistics. For Bayes factors: **bolded** = substantial evidence (>3:1 odds) for a correlational relationship; For *p*-values: *=survives FDR correction for multiple comparisons within each family of tests (families defined by summary statistics)

| Region | Hit MRT |  |  | Hit SDRT |  |  |
| --- | --- | --- | --- | --- | --- | --- |
| | $r$ [CI] | $BF_{10}$ | $p$ | $r$ [CI] | $BF_{10}$ | $p$ |
| CR/FA Left Striatum (-20,16,-2) | -.21[-.36,-.05] | 2.54 | .011* | -.04[-.20,.12] | 0.12 | .626 |
| CR/FA Right Parietal (36,-38,50) | .10[-.06,.26] | 0.22 | .217 | .19[.03,.34] | 1.31 | .024 |
| CR/FA Right Striatum (20,12,-10) | -.22[-.36,-.05] | 2.91 | .010* | -.01[-.17,.15] | 0.11 | .898 |
| CR/GO Left Ant. Insula/IFG (-36,16,6) | -.10[-.26,.07] | 0.20 | .245 | .05[-.12,.21] | 0.12 | .582 |
| CR/GO Left Middle Frontal (-34,38,26) | -.17[-.32,.00] | 0.71 | .049 | -.12[-.28,.04] | 0.29 | .150 |
| CR/GO Left Parietal (-58,-48,34) | -.15[-.30,.01] | 0.52 | .072 | .01[-.15,.18] | 0.11 | .870 |
| CR/GO Right Ant. Insula/IFG (34,20,0) | -.24[-.38,-.07] | **5.60** | .005* | -.08[-.24,.08] | 0.17 | .322 |
| CR/GO Right Middle Frontal (40,42,22) | -.27[-.41,-.11] | **17.98** | .001* | -.23[-.38,-.07] | **4.98** | .005* |
| CR/GO Right Parietal (58,-44,28) | -.33[-.46,-.17] | **255.00** | 7.2E-5* | -.20[-.35,-.03] | 1.57 | .019 |
| CR/GO Supp. Motor Area (8,4,54) | -.14[-.29,.03] | 0.38 | .108 | .01[-.15,.17] | 0.11 | .919 |
| FA/CR Anterior Cingulate (2,30,24) | -.14[-.29,.03] | 0.38 | .107 | -.30[-.44,-.14] | **80.30** | 2.5E-4* |
| FA/CR Left Ant. Insula/IFG (-42,12,-4) | -.05[-.22,.11] | 0.13 | .518 | -.24[-.38,-.07] | **5.48** | .005* |
| FA/CR Left Middle Frontal (-26,48,22) | -.12[-.27,.05] | 0.27 | .171 | -.22[-.37,-.06] | **3.13** | .009* |
| FA/CR Right Ant. Insula/IFG (44,12,-4) | -.04[-.20,.13] | 0.12 | .666 | -.24[-.39,-.08] | **6.84** | .004* |
| FA/CR Right Middle Frontal (24,52,24) | -.05[-.21,.11] | 0.13 | .529 | -.14[-.30,.02] | 0.42 | .095 |
| FA/CR Thalamus_(-14,-16,8) | .09[-.8,.24] | 0.18 | .307 | .00[-.16,.16] | 0.11 | .994 |

| Region | FA rate |  |  |
| --- | --- | --- | --- |
| | $r$ [CI] | $BF_{10}$ | $p$ |
| CR/FA Left Striatum (-20,16,-2) | .19[.03,.34] | 1.51 | .020* |
| CR/FA Right Parietal (36,-38,50) | .03[-.13,.19] | 0.11 | .698 |
| CR/FA Right Striatum (20,12,-10) | .22[.06,.37] | **3.65** | .007* |
| CR/GO Left Ant. Insula/IFG (-36,16,6) | .20[.04,.35] | 1.86 | .016* |
| CR/GO Left Middle Frontal (-34,38,26) | .09[-.08,.25] | 0.18 | .285 |
| CR/GO Left Parietal (-58,-48,34) | .30[.14,.44] | **82.83** | 4.4E-4* |
| CR/GO Right Ant. Insula/IFG (34,20,0) | .24[.07,.38] | **5.74** | .004* |
| CR/GO Right Middle Frontal (40,42,22) | .08[-.09,.24] | 0.16 | .362 |
| CR/GO Right Parietal (58,-44,28) | .11[-.06,.27] | 0.24 | .200 |
| CR/GO Supp. Motor Area (8,4,54) | .07[-.09,.23] | 0.15 | .401 |
| FA/CR Anterior Cingulate (2,30,24) | -.40[-.53,-.25] | **1.9E+4** | 7.7E-7* |
| FA/CR Left Ant. Insula/IFG (-42,12,-4) | -.42[-.54,-.27] | **8.5E+4** | 1.6E-7* |
| FA/CR Left Middle Frontal (-26,48,22) | -.25[-.40,-.09] | **10.48** | .002* |
| FA/CR Right Ant. Insula/IFG (44,12,-4) | -.49[-.60,-.35] | **2.6E+7** | 4.1E-10* |
| FA/CR Right Middle Frontal (24,52,24) | -.40[-.53,-.25] | **1.9E+4** | 7.5E-7* |
| FA/CR Thalamus (-14,-16,8) | -.09[-.25,.07] | 0.19 | .275 |

**Supplemental Table 2a.**
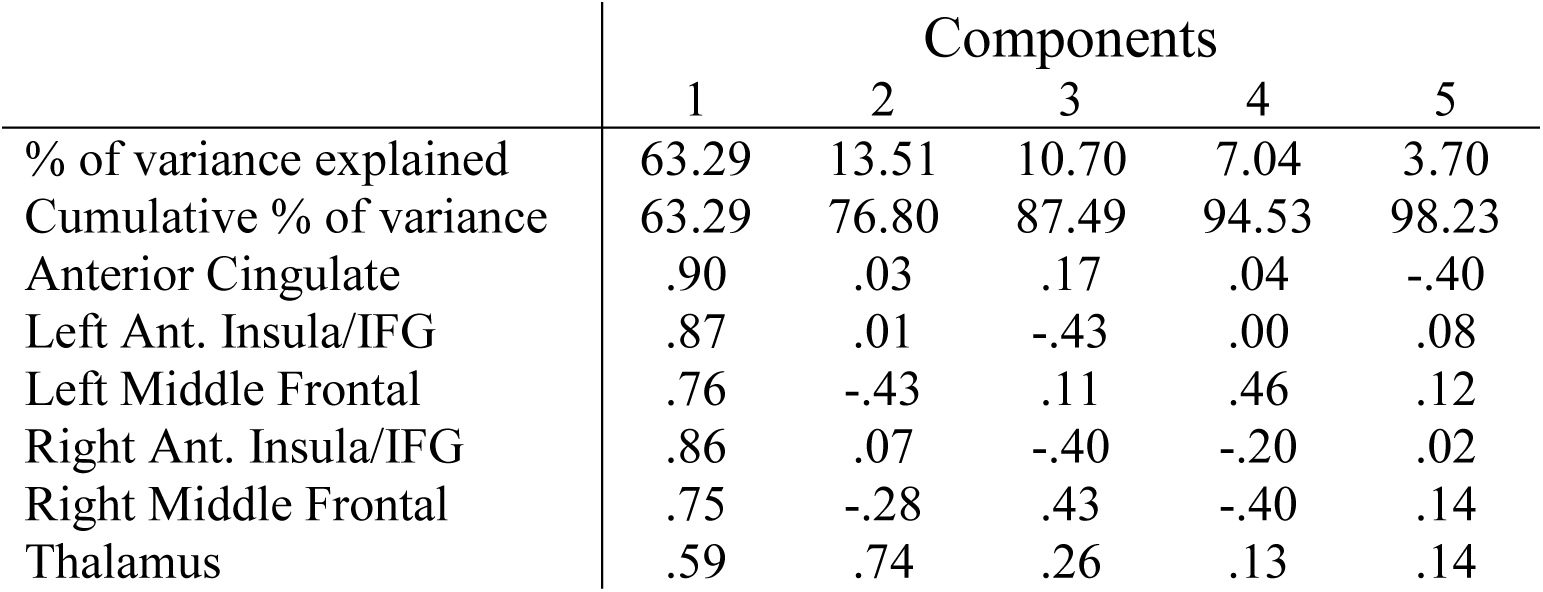
Outcome of the principal component analysis of neural responses in ROIs from the CR>FA contrast. Values in rows labeled with ROI names indicate correlations between each ROI and component.

**Supplemental Table 2a.** Correlation (*r*) values, 95% posterior credible intervals (CIs), Bayes factors (*BF_10_*) and frequentist *p*-values for relationships between model-based indices, including the first component of the FA>CR PCA analysis (“FA>CR.comp”), and externalizing psychopathology self-ratings. All analyses excluded the 5 subjects who did not have ASR data available. CIs and *r* values were estimated assuming a uniform prior from −1 to 1. Bayes factors for *v*.behavior, *v*.brain, *a*, and FA>CR.comp tested the directional hypothesis that the relationship with psychopathology was negative (i.e., the uniform prior spanned values between −1 and 0). Bayes factors for *z* and *Ter* tested the hypothesis that the relationship with psychopathology was positive (i.e., the uniform prior spanned values between 0 and 1). For Bayes factors: **bolded** = substantial evidence (>3:1 odds) for the tested correlational relationship; For *p*-values: *=survives FDR correction for multiple comparisons within each family of tests (families defined by psychopathology measure)

| Predictor | Externalizing Scale (raw) |  |  | DSM-ADHD Scale (raw) |  |  |
| --- | --- | --- | --- | --- | --- | --- |
| | $r$ [CI] | $BF_{10}$ | $p$ | $r$ [CI] | $BF_{10}$ | $p$ |
| $v$.behavior | -.13[-.29,.04] | .64 | .124 | -.21[-.36,-.05] | **4.70** | .012* |
| $v$.brain | -.23[-.38,-.06] | **7.90** | .007* | -.28[-.42,-.12] | **48.29** | 9.3E-04* |
| FA>CR.comp | -.21[-.36,-.04] | **4.04** | .015* | -.25[-.39,-.08] | **14.56** | .003* |
| $a$ | -.07[-.24,.09] | .25 | .393 | -.07[-.23,.10] | .23 | .418 |
| $z$ | -.06[-.22,.11] | .07 | .497 | -.03[-.19,.14] | .08 | .749 |
| $Ter$ | -.03[-.20,.14] | .08 | .716 | .04[-.12,.21] | .17 | .614 |

## References

1. Endres, M. J., Rickert, M. E., Bogg, T., Lucas, J., & Finn, P. R. (2011). Externalizing psychopathology and behavioral disinhibition: Working memory mediates signal discriminability and reinforcement moderates response bias in approach–avoidance learning. Journal of Abnormal Psychology, 120(2), 336.

2. Metin, B., Roeyers, H., Wiersema, J. R., van der Meere, J., & Sonuga-Barke, E. (2012). A meta-analytic study of event rate effects on Go/No-Go performance in attention-deficit/hyperactivity disorder. Biological Psychiatry, 72(12), 990–996.

3. Wright, L., Lipszyc, J., Dupuis, A., Thayapararajah, S. W., & Schachar, R. (2014). Response inhibition and psychopathology: A meta-analysis of go/no-go task performance. Journal of Abnormal Psychology, 123(2), 429.

4. Castellanos-Ryan, N., Struve, M., Whelan, R., Banaschewski, T., Barker, G. J., Bokde, A. L., … & Frouin, V. (2014). Neural and cognitive correlates of the common and specific variance across externalizing problems in young adolescence. American Journal of Psychiatry, 171(12), 1310–1319.

5. Saunders, B., Farag, N., Vincent, A. S., Collins Jr, F. L., Sorocco, K. H., & Lovallo, W. R. (2008). Impulsive errors on a Go-NoGo reaction time task: disinhibitory traits in relation to a family history of alcoholism. Alcoholism: Clinical and Experimental Research, 32(5), 888–894.

6. Verdejo-García, A., Lawrence, A. J., & Clark, L. (2008). Impulsivity as a vulnerability marker for substance-use disorders: review of findings from high-risk research, problem gamblers and genetic association studies. Neuroscience & Biobehavioral Reviews, 32(4), 777–810.

7. Winstanley, C. A., Eagle, D. M., & Robbins, T. W. (2006). Behavioral models of impulsivity in relation to ADHD: translation between clinical and preclinical studies. Clinical Psychology Review, 26(4), 379–395.

8. Smith, J. L., Mattick, R. P., Jamadar, S. D., & Iredale, J. M. (2014). Deficits in behavioural inhibition in substance abuse and addiction: a meta-analysis. Drug and Alcohol Dependence, 145, 1–33.

9. Casey, B. J., Trainor, R. J., Orendi, J. L., Schubert, A. B., Nystrom, L. E., Giedd, J. N., … & Forman, S. D. (1997). A developmental functional MRI study of prefrontal activation during performance of a go-no-go task. Journal of Cognitive Neuroscience, 9(6), 835–847.

10. Durston, S., Thomas, K. M., Yang, Y., Uluğ, A. M., Zimmerman, R. D., & Casey, B. J. (2002). A neural basis for the development of inhibitory control. Developmental Science, 5(4), F9–F16.

11. Durston, S., Tottenham, N. T., Thomas, K. M., Davidson, M. C., Eigsti, I. M., Yang, Y., … & Casey, B. J. (2003). Differential patterns of striatal activation in young children with and without ADHD. Biological Psychiatry, 53(10), 871–878.

12. Norman, A. L., Pulido, C., Squeglia, L. M., Spadoni, A. D., Paulus, M. P., & Tapert, S. F. (2011). Neural activation during inhibition predicts initiation of substance use in adolescence. Drug and Alcohol Dependence, 119(3), 216–223.

13. Schulz, K. P., Fan, J., Tang, C. Y., Newcorn, J. H., Buchsbaum, M. S., Cheung, A. M., & Halperin, J. M. (2004). Response inhibition in adolescents diagnosed with attention deficit hyperactivity disorder during childhood: an event-related FMRI study. American Journal of Psychiatry, 161(9), 1650–1657.

14. Wetherill, R. R., Squeglia, L. M., Yang, T. T., & Tapert, S. F. (2013). A longitudinal examination of adolescent response inhibition: neural differences before and after the initiation of heavy drinking. Psychopharmacology, 230(4), 663–671.

15. Garavan, H., Ross, T. J., & Stein, E. A. (1999). Right hemispheric dominance of inhibitory control: an event-related functional MRI study. Proceedings of the National Academy of Sciences, 96(14), 8301–8306.

16. Zheng, D., Oka, T., Bokura, H., & Yamaguchi, S. (2008). The key locus of common response inhibition network for no-go and stop signals. Journal of Cognitive Neuroscience, 20(8), 1434–1442.

17. Criaud, M., & Boulinguez, P. (2013). Have we been asking the right questions when assessing response inhibition in go/no-go tasks with fMRI? A meta-analysis and critical review. Neuroscience & Biobehavioral Reviews, 37(1), 11–23.

18. Swick, D., Ashley, V., & Turken, U. (2011). Are the neural correlates of stopping and not going identical? Quantitative meta-analysis of two response inhibition tasks. Neuroimage, 56(3), 1655–1665.

19. Ahmadi, A., Pearlson, G. D., Meda, S. A., Dager, A., Potenza, M. N., Rosen, R., … & Wood, R. M. (2013). Influence of alcohol use on neural response to go/no-go task in college drinkers. Neuropsychopharmacology, 38(11), 2197.

20. Claus, E. D., Feldstein Ewing, S. W., Filbey, F. M., & Hutchison, K. E. (2013). Behavioral control in alcohol use disorders: relationships with severity. Journal of Studies on Alcohol and Drugs, 74(1), 141–151.

21. Garavan, H., Ross, T. J., Murphy, K., Roche, R. A. P., & Stein, E. A. (2002). Dissociable executive functions in the dynamic control of behavior: inhibition, error detection, and correction. Neuroimage, 17(4), 1820–1829.

22. Stevens, M. C., Kiehl, K. A., Pearlson, G. D., & Calhoun, V. D. (2009). Brain network dynamics during error commission. Human Brain Mapping, 30(1), 24–37.

23. Heitzeg, M. M., Nigg, J. T., Hardee, J. E., Soules, M., Steinberg, D., Zubieta, J. K., & Zucker, R. A. (2014). Left middle frontal gyrus response to inhibitory errors in children prospectively predicts early problem substance use. Drug and Alcohol Dependence, 141, 51–57.

24. Huster, R. J., Eichele, T., Enriquez-Geppert, S., Wollbrink, A., Kugel, H., Konrad, C., & Pantev, C. (2011). Multimodal imaging of functional networks and event-related potentials in performance monitoring. Neuroimage, 56(3), 1588–1597.

25. Rasmussen, J., Casey, B. J., van Erp, T. G., Tamm, L., Epstein, J. N., Buss, C., … & Somerville, L. (2016). ADHD and cannabis use in young adults examined using fMRI of a Go/NoGo task. Brain Imaging and Behavior, 10(3), 761–771.

26. Czapla, M., Baeuchl, C., Simon, J. J., Richter, B., Kluge, M., Friederich, H. C., … & Loeber, S. (2017). Do alcohol-dependent patients show different neural activation during response inhibition than healthy controls in an alcohol-related fMRI go/no-go-task?. Psychopharmacology, 234(6), 1001–1015.

27. Dillo, W., Göke, A., Prox-Vagedes, V., Szycik, G. R., Roy, M., Donnerstag, F., … & Ohlmeier, M. D. (2010). Neuronal correlates of ADHD in adults with evidence for compensation strategies–a functional MRI study with a Go/No-Go paradigm. GMS German Medical Science, 8.

28. Ding, W. N., Sun, J. H., Sun, Y. W., Chen, X., Zhou, Y., Zhuang, Z. G., … & Du, Y. S. (2014). Trait impulsivity and impaired prefrontal impulse inhibition function in adolescents with internet gaming addiction revealed by a Go/No-Go fMRI study. Behavioral and Brain Functions, 10(1), 20.

29. Tapert, S. F., Schweinsburg, A. D., Drummond, S. P., Paulus, M. P., Brown, S. A., Yang, T. T., & Frank, L. R. (2007). Functional MRI of inhibitory processing in abstinent adolescent marijuana users. Psychopharmacology, 194(2), 173–183.

30. Ratcliff, R. (1978). A theory of memory retrieval. Psychological Review, 85(2), 59.

31. Ratcliff, R., Smith, P. L., Brown, S. D., & McKoon, G. (2016). Diffusion decision model: current issues and history. Trends in Cognitive Sciences, 20(4), 260–281.

32. Ratcliff, R., Huang-Pollock, C., & McKoon, G. (2018). Modeling individual differences in the go/no-go task with a diffusion model. Decision, 5(1), 42.

33. Huang-Pollock, C., Ratcliff, R., McKoon, G., Shapiro, Z., Weigard, A., & Galloway-Long, H. (2017). Using the diffusion model to explain cognitive deficits in attention deficit hyperactivity disorder. Journal of Abnormal Child Psychology, 45(1), 57–68.

34. Zucker, R. A., Ellis, D. A., Fitzgerald, H. E., Bingham, C. R., & Sanford, K. (1996). Other evidence for at least two alcoholisms II: Life course variation in antisociality and heterogeneity of alcoholic outcome. Development and Psychopathology.

35. Zucker, R. A., Fitzgerald, H. E., Refior, S. K., Puttler, L. I., Pallas, D. M., & Ellis, D. A. (2000). The clinical and social ecology of childhood for children of alcoholics: Description of a study and implications for a differentiated social policy. In: Fitzgerald, H.E., Lester, B.M., & Zucker, R.A. (Eds), Children of Addiction: Research, Health and Policy Issues. (pp. 109–141). New York: Routledge Falmer Publishers.

36. Glover, G. H., & Law, C. S. (2001). Spiral-in/out BOLD fMRI for increased SNR and reduced susceptibility artifacts. Magnetic Resonance in Medicine, 46(3), 515–522.

37. Achenbach, T. M., & Rescorla, L. (2003). Manual for the ASEBA adult forms & profiles: for ages 18-59: adult self-report and adult behavior checklist. ASEBA.

38. R Core Team (2018). R: A Language and Environment for Statistical Computing, R Foundation for Statistical Computing, Austria, 2015.

39. Singmann, H., Brown, S., Gretton, M., & Heathcote, A. (2016). rtdists: Response time distributions. R package version 0.4-9. URL http://CRAN.R-project.org/package=rtdists.

40. Voss, A., Nagler, M., & Lerche, V. (2013). diffusion Models in Experimental Psychology: A Practical Introduction. Experimental Psychology, 60(6), 385–402.

41. Dutilh, G., Annis, J., Brown, S. D., Cassey, P., Evans, N. J., Grasman, R. P., … & Kupitz, N. (2016). The quality of response time data inference: A blinded, collaborative assessment of the validity of cognitive models. Psychonomic Bulletin & Review, 1–19.

42. Fessler, J. A., Lee, S., Olafsson, V. T., Shi, H. R., & Noll, D. C. (2005). Toeplitz-based iterative image reconstruction for MRI with correction for magnetic field inhomogeneity. IEEE Transactions on Signal Processing, 53(9), 3393–3402.

43. Brett, M., Anton, J. L., Valabregue, R., & Poline, J. B. (2002, June). Region of interest analysis using an SPM toolbox. In 8th international conference on functional mapping of the human brain (Vol. 16, No. 2, p. 497).

44. JASP Team. (2018). JASP (Version 0.9. 0.1)[Computer software].

45. Dienes, Z. (2016). How Bayes factors change scientific practice. Journal of Mathematical Psychology, 72, 78–89.

46. Wagenmakers, E. J., Marsman, M., Jamil, T., Ly, A., Verhagen, J., Love, J., … & Matzke, (2018). Bayesian inference for psychology. Part I: Theoretical advantages and practical ramifications. Psychonomic Bulletin & Review, 25(1), 35–57.

47. Benjamini, Y., & Hochberg, Y. (1995). Controlling the false discovery rate: a practical and powerful approach to multiple testing. Journal of the Royal Statistical Society: Series B (Methodological), 57(1), 289–300.

48. Rosseel, Y., Oberski, D., Byrnes, J., Vanbrabant, L., Savalei, V., Merkle, E., … & Chow, M. (2018). Package ‘lavaan’.

49. Martz, M. E., Zucker, R. A., Schulenberg, J. E., & Heitzeg, M. M. (2018). Psychosocial and neural indicators of resilience among youth with a family history of substance use disorder. Drug and Alcohol Dependence, 185, 198–206.

50. Metin, B., Roeyers, H., Wiersema, J. R., van der Meere, J. J., Thompson, M., & Sonuga-Barke, E. (2013). ADHD performance reflects inefficient but not impulsive information processing: A diffusion model analysis. Neuropsychology, 27(2), 193.

51. Weigard, A., & Huang-Pollock, C. (2017). The role of speed in ADHD-related working memory deficits: a time-based resource-sharing and diffusion model account. Clinical Psychological Science, 5(2), 195–211.

52. Ziegler, S., Pedersen, M. L., Mowinckel, A. M., & Biele, G. (2016). Modelling ADHD: a review of ADHD theories through their predictions for computational models of decision-making and reinforcement learning. Neuroscience & Biobehavioral Reviews, 71, 633–656.

53. Chambers, C. D., Garavan, H., & Bellgrove, M. A. (2009). Insights into the neural basis of response inhibition from cognitive and clinical neuroscience. Neuroscience & Biobehavioral Reviews, 33(5), 631–646.

54. Stevens, M. C., Kiehl, K. A., Pearlson, G. D., & Calhoun, V. D. (2007). Functional neural networks underlying response inhibition in adolescents and adults. Behavioural Brain Research, 181(1), 12–22.

55. Endres, M. J., Donkin, C., & Finn, P. R. (2014). An information processing/associative learning account of behavioral disinhibition in externalizing psychopathology. Experimental and Clinical Psychopharmacology, 22(2), 122.

56. Bohlin, G., Eninger, L., Brocki, K. C., & Thorell, L. B. (2012). Disorganized attachment and inhibitory capacity: Predicting externalizing problem behaviors. Journal of Abnormal Child Psychology, 40(3), 449–458.

57. Barkley, R. A. (1997). Behavioral inhibition, sustained attention, and executive functions: constructing a unifying theory of ADHD. Psychological Bulletin, 121(1), 65.

58. Karalunas, S. L., Geurts, H. M., Konrad, K., Bender, S., & Nigg, J. T. (2014). Annual research review: Reaction time variability in ADHD and autism spectrum disorders: Measurement and mechanisms of a proposed trans-diagnostic phenotype. Journal of Child Psychology and Psychiatry, 55(6), 685–710.

59. Weigard, A., Huang-Pollock, C., Brown, S., & Heathcote, A. (2018). Testing formal predictions of neuroscientific theories of ADHD with a cognitive model–based approach. Journal of Abnormal Psychology, 127(5), 529.

60. Aston-Jones, G., & Cohen, J. D. (2005). An integrative theory of locus coeruleus-norepinephrine function: adaptive gain and optimal performance. Annual Review of Neuroscience, 28, 403–450.

## Supplemental References

1. Glover, G. H., & Law, C. S. (2001). Spiral-in/out BOLD fMRI for increased SNR and reduced susceptibility artifacts. Magnetic Resonance in Medicine, 46(3), 515–522.

2. Ratcliff, R., & Tuerlinckx, F. (2002). Estimating parameters of the diffusion model: Approaches to dealing with contaminant reaction times and parameter variability. Psychonomic Bulletin & Review, 9(3), 438–481.

3. Voss, A., Nagler, M., & Lerche, V. (2013). Diffusion models in experimental psychology. Experimental Psychology.

4. Dutilh, G., Annis, J., Brown, S. D., Cassey, P., Evans, N. J., Grasman, R. P., … & Kupitz, C. N. (2016). The quality of response time data inference: A blinded, collaborative assessment of the validity of cognitive models. Psychonomic Bulletin & Review, 1–19.

5. Ratcliff, R., Huang-Pollock, C., & McKoon, G. (2018). Modeling individual differences in the go/no-go task with a diffusion model. Decision, 5(1), 42.

6. Huang-Pollock, C., Ratcliff, R., McKoon, G., Shapiro, Z., Weigard, A., & Galloway-Long, H. (2017). Using the diffusion model to explain cognitive deficits in attention deficit hyperactivity disorder. Journal of Abnormal Child Psychology, 45(1), 57–68.

7. Singmann, H., Brown, S., Gretton, M., & Heathcote, A. (2016). rtdists: Response time distributions. R package version 0.4-9. URL http://CRAN.R-project.org/package=rtdists.

8. Voss, A., & Voss, J. (2007). Fast-dm: A free program for efficient diffusion model analysis. Behavior Research Methods, 39(4), 767–775.

9. Wagenmakers, E. J., Van Der Maas, H. L., & Grasman, R. P. (2007). An EZ-diffusion model for response time and accuracy. Psychonomic Bulletin & Review, 14(1), 3–22.

10. White, C. N., Servant, M., & Logan, G. D. (2018). Testing the validity of conflict drift-diffusion models for use in estimating cognitive processes: A parameter-recovery study. Psychonomic Bulletin & Review, 25(1), 286–301.

11. Husson, F., Josse, J., Le, S., Mazet, J., & Husson, M. F. (2018). Package ‘FactoMineR’. Package FactorMineR.

